# N-linked glycosylation of the antagonist Short gastrulation increases the functional complexity of BMP signals

**DOI:** 10.1101/316448

**Authors:** E. Negreiros, S. Herszterg, K. Hwa, A. Câmara, W.B. Dias, K. Carneiro, E. Bier, A. Todeschini, H. Araujo

## Abstract

Disorders of N-linked glycosylation are increasingly reported in the literature. However, targets responsible for the associated developmental and physiological defects are largely unknown. Bone Morphogenetic Proteins (BMPs) act as highly dynamic complexes to regulate several functions during development. The range and strength of BMP activity depend on interactions with glycosylated protein complexes in the extracellular milieu. Here we investigate the role of glycosylation for the function of the conserved extracellular BMP antagonist Short gastrulation (Sog). We identify conserved N-glycosylated sites and describe the effect of mutating these residues on BMP pathway activity in Drosophila. Functional analysis reveals that loss of individual Sog glycosylation sites enhances BMP antagonism and/or increases the spatial range of Sog effects in the tissue. Mechanistically, we provide evidence that N-terminal and stem glycosylation controls extracellular Sog levels and distribution. The identification of similar residues in vertebrate Chordin proteins suggests that N-glycosylation may be an evolutionarily conserved process that adds complexity to the regulation of BMP activity.

**Summary Statement:** N-glycosylation restricts the function of Short gastrulation during Drosophila development by controlling the amount of extracellular protein. This adds another layer of complexity to regulation of Bone Morphogenetic Protein signals.

## Introduction

Glycosylation of secreted proteins in multicellular organisms increases structural diversity extending the possibilities of cell responses to the microenvironment. Protein glycosylation depends on a series of reactions taking place in different cellular compartments. Activated nucleotide-sugar precursors are synthetized in the cytoplasm, and imported into the lumen of the endoplasmic reticulum (ER) and Golgi compartments, where these sugar substrates are used as the building blocks. N-linked glycosylation is initiated in the ER by adding a conserved oligosaccharide en bloc from a dolichol-linked precursor oligosaccharide onto newly translated proteins. After an initial trimming in the ER the oligosaccharide chain is processed and modified in the Golgi apparatus (Stanley et al., 2017). A series of glycosyltransferases and multiprotein complexes regulate these steps. Improper sugar trimming is linked to errors in protein folding and protein degradation in the ER. Notably, a series of congenital disorders of glycosylation have been reported, with the majority affecting primarily N-glycan assembly (Freeze et al., 2014).

A great challenge is to understand how glycosylation modulates the function of their molecular targets. Critical targets of glycosylation are the Bone Morphogenetic Proteins (BMPs) and their binding partners, which control embryonic development and tissue patterning (Bier and De Robertis, 2015). Extracellular proteoglycans and glycoproteins modulate BMP signaling in several species. For instance, *Xenopus* and mammalian N-acetylgalactosaminyltransferases inhibit BMP signaling (Herr et al., 2008). In zebrafish, embryos injected with morpholinos against beta1,4- galactosyltransferase have reduced activation of the BMP-dependent transcription factors Smad1/5/8 (Machingo et al., 2006). Likewise, mutation of glycosylation sites in mouse Twisted gastrulation (Tsg), an important regulator of BMP activity, reduces its binding to BMPs (Billington et al., 2011). Glycosylation also controls the secretion and folding of human BMP-2 (Hang et al., 2014), BMP receptor recognition, specificity, and binding strength (Lowery et al., 2014, Saremba et al., 2008).

In Drosophila, activity of the BMPs Decapentaplegic (Dpp), Glass bottom boat (Gbb) and Screw (Scw) depend on glycan-based interactions. Early work has shown that genes involved in proteoglycan biosynthesis such as *division abnormally delayed* (*dally*), *sugarless* (*sgl*), and *sulfateless* (*sfl*) regulate Dpp function (Häcker et al., 2005, Selleck, 2000). Furthermore, glycosylation can transform Drosophila Dpp from a long-range to a short-range signal (Humphreys et al., 2013). However, the role of N-linked glycosylation on Drosophila BMP function is less explored. N-linked glycans in Drosophila are less complex than in vertebrates (North et al., 2006, Aoki et al., 2007, Gagneux and Varki, 1999). Nonetheless, several enzymes in the N-glycosylation pathway have been reported to impair Drosophila development (Wandall et al., 2003, Tian and Ten Hagen, 2006, Yamamoto-Hino et al., 2015).

Dpp activity depends on the formation of protein complexes with Tsg proteins, Tolloid (Tld) metalloproteases and the dedicated antagonist Short gastrulation (Sog). Complex formation is required to regulate Dpp activity range during formation of the veins in the pupal wing and to specify the amnioserosa, the dorsal-most extra-embryonic tissue in the embryo. In the pupal wing, Dpp and Gbb ligands are transported from the longitudinal veins in a complex with Sog, the Tsg-family protein Crossveinless and the Tolloid-related (Tlr) metalloprotease for signaling in the posterior crossvein (PCV) forming area (Serpe et al., 2005, Serpe et al., 2008, Ray and Wharton, 2001). Longitudinal veins also require Sog and Dpp antagonism (Yu et al., 1996) and may depend on shuttling peak amounts of BMPs to the center of the provein domains by the action of Sog, which is produced by the intervein cells (Araujo et al., 2003, Negreiros et al., 2010). During embryogenesis, laterally secreted Sog binds Dpp and Scw in a complex with Tsg that Inhibits Dpp locally and facilitates long-range ligand diffusion towards the dorsal midline. Away from the Sog source, Tld cleaves Sog, delivering peak BMP levels for receptor activation and dorsal amnioserosa formation (Peluso et al., 2011, Sawala et al., 2012, Mizutani et al., 2006, Umulis et al., 2006). Sog is a N-glycosylated protein that presents evolutionarily conserved structure and function (François and Bier, 1995, Marqués et al., 1997). Sog and the vertebrate Chordin homologs regulate dorsal-ventral patterning across phyla by antagonizing BMP activity and regulating BMP spread (Bier and De Robertis, 2015). Furthermore, Sog (Srinivasan et al., 2002) and Chd (Plouhinec et al., 2013) form morphogen gradients in the extracellular space in invertebrate and vertebrate embryos respectively. Since Sog ultimately regulates the pattern of BMP activity in tissues, modifications that alter Sog binding and distribution will potentially have great impact on BMP function. Here we investigate whether Sog glycosylation is important for Sog function. We show that Sog glycosylation mutants alter BMP activity in cells, wings and embryos, consistent with a role in regulating BMP activity during Drosophila development.

## Results

### BMP signaling is impaired in *frc-* wings

*fringe connection* (*frc*) encodes a UDP-sugar transporter that transports a broad range of UDP-sugars for the synthesis of glycans, including N-linked types, GAGs, and mucins (Selva et al., 2001, Goto et al., 2001). *frc* is thus at the basis of glycoprotein and glucosaminoglycan synthesis. Loss-of-function *frc* alleles are pupal lethal and *frc*-wings of adult escapers show notches at the margin, due to impairment of the Notch pathway (Goto et al., 2001). *frc*- wings also display veins of uneven width which may result from the impairment of other signaling pathways as well. Given the implication of the BMP pathway in wing vein patterning, we investigated whether *frc* regulates BMP pathway activity.

During pupal wing development, *dpp* is expressed in the center of broad and irregular (4-7 cells wide) longitudinal vein competent, or provein domains (Blair, 2007). Dpp is required for vein fate maintenance and refinement, and formation of the vein proper requires BMP activity to be restricted to the center of the proveins by the action of both its receptor *thickveins* (*tkv)* and *sog* (de Celis, 1997, Yu et al., 1996). Sog is expressed in the adjacent intervein domains from where it diffuses into the provein territory, likely to control the transport of Dpp outwards and along the veins (Yu et al., 1996, Araujo et al., 2003, Negreiros et al., 2010). BMP activity in the provein domains can be detected with anti-phosphorylated Mad (pMad) antisera. From 24 hours after puparium formation (h APF) to wing expansion, pMad becomes restricted to quasi-continuous 3-4 cells wide domains along the longitudinal veins, as well as in the posterior cross vein (PCV) (Fig. 1D) (de Celis, 1997, Conley et al., 2000). We observed that, in *frc-* wings, pMad staining is patchy along longitudinal veins and enlarged at the PCV, consistent with the resulting veins of uneven width and extra PCV tissue in adult wings (Fig. 1C). This pattern could result from uneven BMP diffusion and/or uneven BMP receptor activation within the provein domain, based on modified interactions with glycosylated molecules.

**Figure 1:**
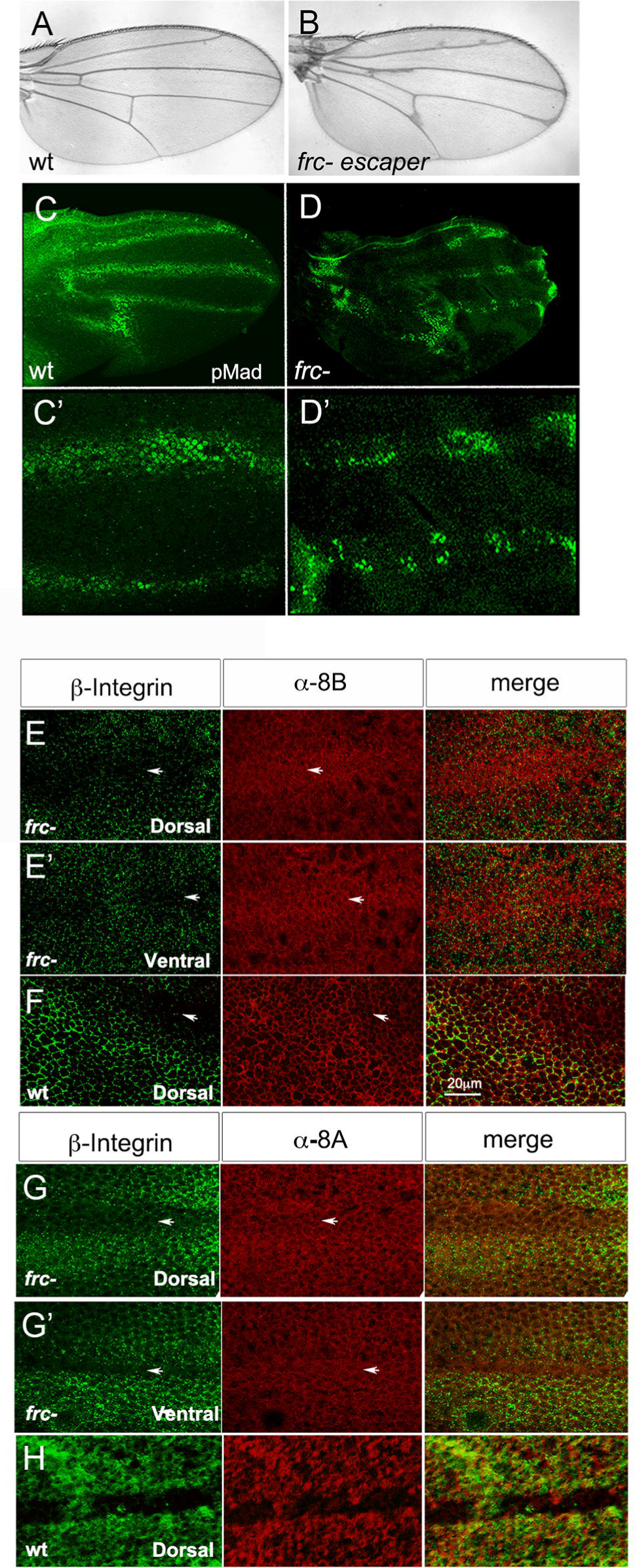
*frc* mutants affect BMP activity and modify Sog distribution in the pupal wing. A) Wild type and B) *frc-*adult wing. C,D) Wild type (C,C’) and *frc-* (D,D-) 24h apf wings stained for pMAD. High magnification shows continuity of pMAD staining in wild type (C’) and a patchy pattern in *frc-* provein staining (D’). E-H) Double immunolabelling for β-integrin and anti-8B (E,F) or anti-8A (J,H) Sog antisera in *frc-* (E,G) or wild type (F,H) 24h apf wings. Shown are the dorsal (E,F,G,H) and ventral (E’,G’) wing epithelia. Note Sog staining inside the provein territory in both the dorsal and ventral wing epithelia in *frc-*.

Sog distribution is also altered in *frc-* wings. We have shown that Sog diffusion from intervein-producing cells into the provein domains takes place only at the dorsal surface of wild-type wings and is restricted to the basolateral domain of the dorsal epithelium (Negreiros et al., 2010). Furthermore, the use of antisera against N- and C-terminal Sog epitopes (8A and 8B, respectively) suggests that only C-terminal Sog fragments are able to enter the provein domain (Araujo et al., 2003, Negreiros et al., 2010) (Figs 1G and 1I). However, in *frc-* wings, both the N- and C-terminal epitopes of Sog are detected in the provein domains (Fig. 1F,H). These results suggest that glycosylation is necessary to control the distribution of Sog between the intervein and vein-competent domains and that Sog interactions based on glycosylated residues could be modified in *frc-* mutants. Since *frc* modifies the loss-of-vein phenotype of enhancer piracy *sog* lines (Fig. S1) and Sog is a glycoprotein (Marqués et al., 1997), we decided to investigate whether Sog glycosylation regulates BMP function.

### Sog and Chordin putative glycosylation sites are conserved

It has been known for a long time that Sog is a glycoprotein (Marqués et al., 1997). However, how glycosylation contributes to Sog function has not been investigated. The evolutionary conservation of putative glycosylation sites suggests a positive selection for these sites in protostome and deuterostome lineages. We have aligned Sog sequences from the 12 sequenced Drosophila species and found that, among the six predicted glycosylation sites, three are conserved in all species analyzed (Fig. 2A; Fig. S2B). The first conserved site is located after the first cystein rich (CR) domain, in close proximity to a Tld cleavage site (Peluso et al., 2011, Marqués et al., 1997) (Fig. 2B). The second and third sites lie in the stem and after the second CR domain, respectively (Fig. 2B). Putative glycosylation sites in locations corresponding to the second site are consistently detected in vertebrate species as well (Fig. S2B).

**Figure 2:**
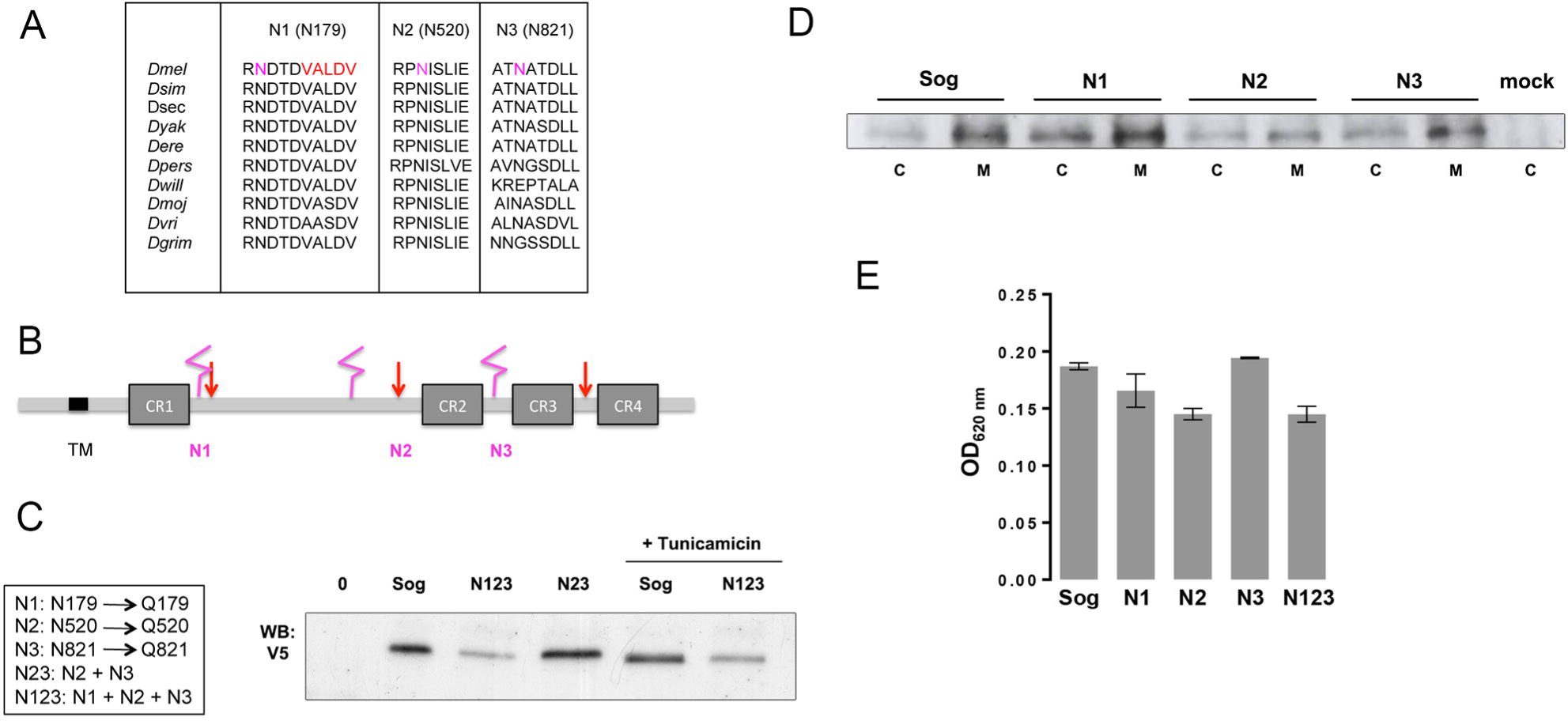
*Drosophila sog* has three conserved glycosylation sites. A) Alignment of Sog sequences from *Drosophila sp*. shows that three putative glycosylation sites are conserved (Arg residue in pink in *D. melanogaster*). In red a Tld/Tlr cleavage site sequence close to N1 is shown. B) Sog protein scheme with the location of the four conserved Cystein Rich (CR) domains, Tld/Tlr cleavage sites (red arrows) and putative glycosylation sites (pink). C-E) Analysis of Sog constructs expressed in S2 cells. C) Glycosylation mutants produced by site-directed mutagenesis and their effect on Sog migration in SDS-PAGE. N23 and N123 migrate faster that wild type Sog. All mutants and wild type Sog bear a C-terminal V5/His tag. Treatment with Tunicamycin decreases wild type Sog Mw and confirms that Sog is glycosylated. D) SDS-PAGE for Sog protein in cells (C) and extracellular medium (M) shows that Sog is secreted and that only the N2 mutation slightly decreases Sog secretion. E) Elisa for S2 cell secreted Sog confirms the analysis in D.

To investigate a functional role for Sog glycosylation, we generated single, double and triple Sog glycosylation mutant constructs, in which we abolished the conserved putative N-glycosylation sites (Asn-X-Ser/Thr) by mutating the Asn residues to Gln. We refer to these constructs as SogN mutants (N1, N2, N3, N23 and N123) (Fig. 2C). Expression of V5/His-tagged double and triple mutants in S2 cells revealed that these mutants have a reduced molecular weight when compared to wild-type Sog, as seen on SDS-PAGE (Fig. 2C). Treatment of cells with tunicamycin, a specific inhibitor of N-linked glycosylation, abolishes this difference (Fig. 2C). This strongly suggests that the identified sites are N-glycosylated and that a decrease in the number of sugar side chains is responsible for the differential migration pattern of SogN mutants.

Since loss of glycosylation is frequently associated with impaired protein folding (Caramelo and Parodi, 2015) we tested whether SogN mutants are appropriately secreted. We transfected S2 cells with equivalent amounts of wild type or mutated epitope tagged *sog* constructs. After 48h of induction, we collected cell (C) or medium (M) samples for Western blot and Elisa. Secretion of Sog into the medium was slightly decreased only for constructs bearing the second mutated site (N2 and N123) and was unaffected for the other mutants (Fig. 2D,E). Therefore, eventual phenotypes resulting from *sogN* expression (see below) are unlikely to result from impaired protein secretion.

### Sog glycosylation mutants display increased BMP binding

Next, we analyzed whether SogN mutants displayed differences in BMP binding properties and cleavage by metalloproteases. Full-length Sog binds both Dpp homodimers and heterodimers, but shows a greater affinity for Dpp heterodimers (Fig. 3A, compare lanes 2 and 4) (Shimmi et al., 2005). This preference is modified by the presence of Tsg, which increases the affinity of full-length Sog for Dpp homodimers (Fig. 3A, lanes 5 and 6). This behavior is distinct for the N-terminal Supersog cleavage product, which is able to bind Dpp alone (Fig. 3B, lanes 4 and 6). Interestingly, the Sog triple glycosylation mutant N123 shows increased binding to Dpp homodimers in the absence of Tsg (Fig. 3C), suggesting that Sog glycosylation modulates Sog binding to BMPs. Among the single glycosylation mutants, SogN1 was the sole mutant showing enhanced binding to Dpp homodimers in the absence of another BMP or Tsg (Fig. 3D, lane 7).

**Figure 3:**
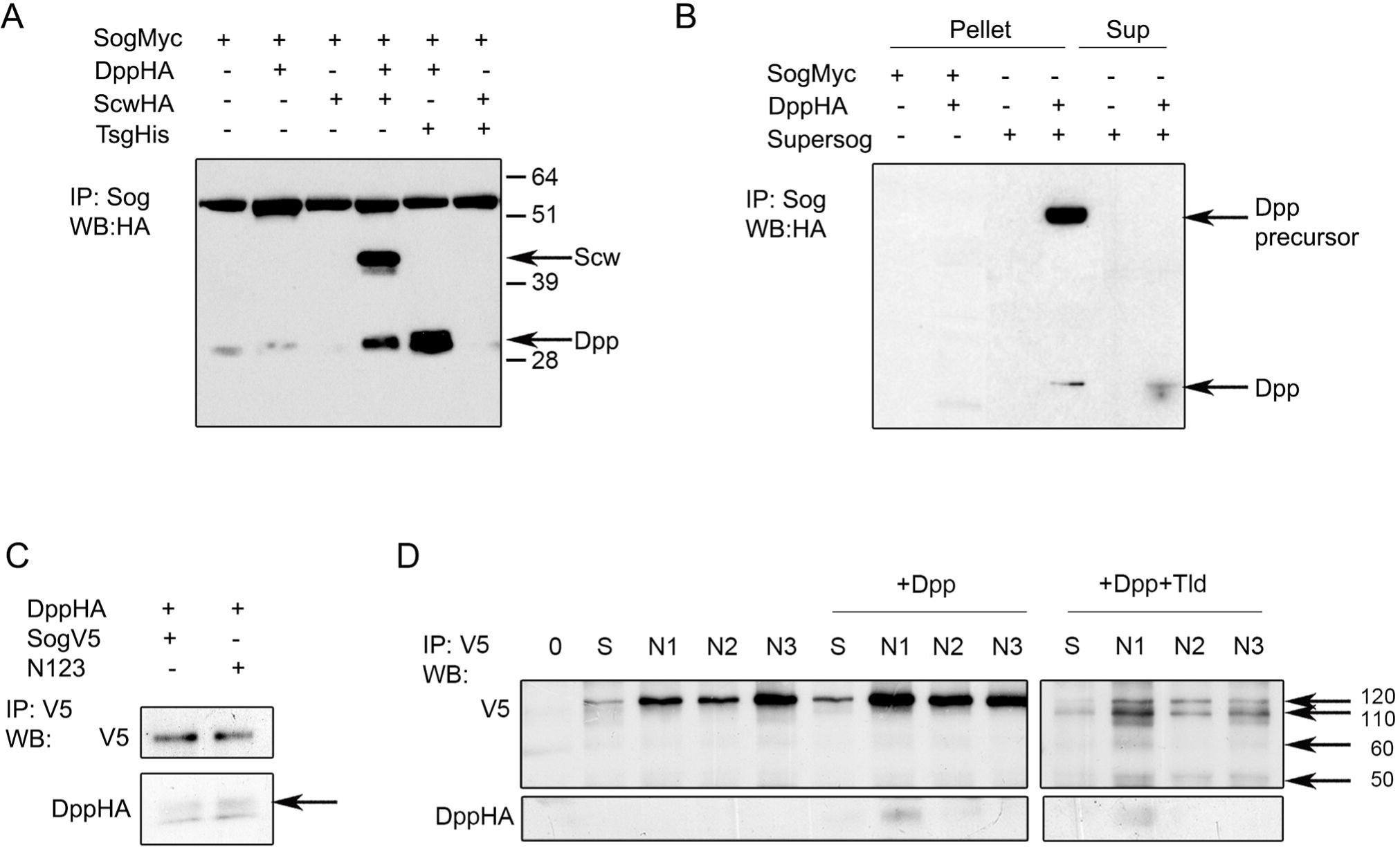
Loss of Sog glycosylation modifies Dpp binding. A) S2 cells transfected with wild type Sog-myc and different combinations of Tsg and BMP ligands. Co-IP with cell supernatants shows that Sog binds only BMP heterodimers in the absence of Tsg, but binds Dpp homodimers in the presence of Tsg. B) S2 cells transfected with wild type Sog-myc or the N-terminal Supersog fragment reveal that Supersog binds Dpp alone, either using cell pellets or supernatants. C) S2 cells transfected with wild type *sog-*V5 or *sogN123*-V5, and *dpp-*HA. Co-IP for V5 shows that the N123 mutant binds more Dpp than wild type Sog. D) S2 cells transfected with wild type *sog* (S) or single glycosylation mutants (N1, N2, N3) plus *dpp*-HA or *dpp-*HA and *tld*-HA. Co-IP for V5. SogN1 binds Dpp alone, while other mutants and wild type Sog do not. Arrows point to the different Sog fragments produced by the Tld metalloprotease and show that the cleavage pattern is similar among all constructs.

Sog glycosylation could also affect Sog cleavage by Tld metalloproteases and impact on the consequent release of BMPs for receptor binding. Unlike vertebrate Chordin, Sog cleavage by Tld metalloproteases relies on the presence of BMPs (Marqués et al., 1997, Peluso et al., 2011). We found that SogN mutants display a cleavage pattern similar to wild type Sog (Fig. 3D, right panel and Yu et al., 2000). Therefore, loss of Sog glycosylation does not affect cleavage by Tld metalloproteases, although a decrease in cleavage efficiency cannot be ruled out in these experiments.

### Loss of glycosylation sites enhances Sog function *in vivo*

The differences in BMP binding described above suggested that loss of glycosylation could modify Sog function *in vivo*. To test this prediction, we assayed the effects of overexpressing *sog* glycosylation mutants during embryogenesis and pupal wing development. During pupal wing development, Dpp is expressed in the longitudinal veins where it establishes the vein proper domain (de Celis, 1997, Sotillos and De Celis, 2005). In addition, Dpp is transported from longitudinal veins to initiate BMP signals and formation of the PCV (Conley et al., 2000, Serpe et al., 2005, Ralston and Blair, 2005, Shimmi et al., 2005)Shimmi et al., 2005). Ubiquitous overexpression of wild-type *sog* in the wing leads to various degrees of longitudinal vein truncation and PCV loss (Yu et al., 1996, Serpe et al., 2005)(Figure S3 and Figure 4B). Vein-restricted *sog* overexpression by using the shortvein Gal4 driver (*shv*-Gal4) (Sotillos and De Celis, 2005) also results in longitudinal vein truncation to various degrees, as well as partial or complete PCV loss (Fig. 4A,C). We observed that overexpression of *sogN* mutants leads to more severe longitudinal vein truncation phenotypes when compared to wild-type *sog*, with *sogN1* and *sogN2* showing the greatest effect (Fig. 4A). Consistent with inhibition of the BMP pathway exerted by *sog*, overexpression of *sogWT* and *sogN* also increases the severity of longitudinal vein truncation in the *dpp*[shv] background (Fig. S4).

**Figure 4.**
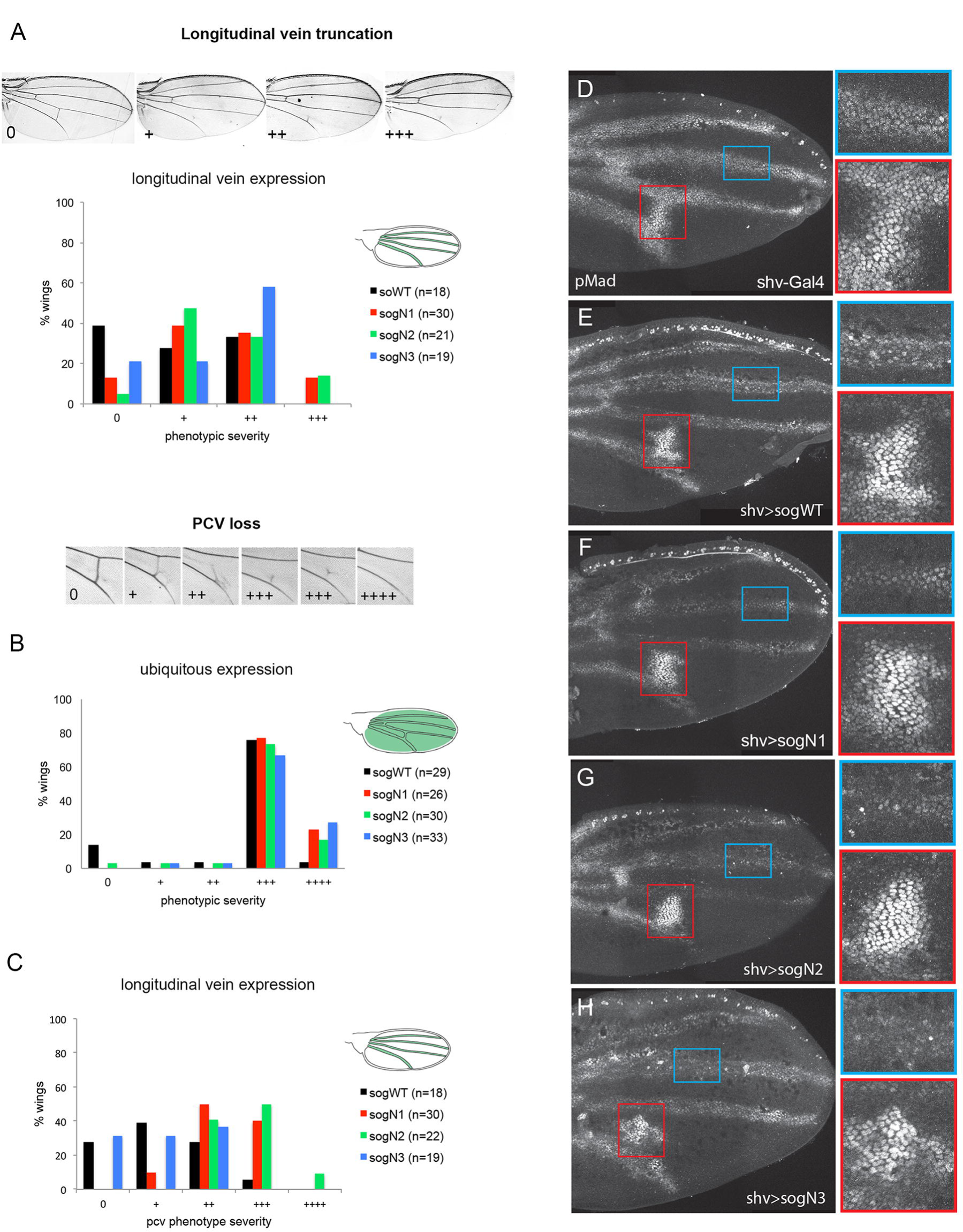
Loss of glycosylation sites enhances Sog function in the wing. A-C) The adult wing venation pattern produced by Gal4/UAS expression of wild type Sog or Sog glycosylation mutants was quantified. Expression induced by the shv-Gal4 driver in the longitudinal veins (A,C) or ubiquitous expression in the wing with the MS1096-Gal4 driver (B) leads to different degrees of longitudinal vein loss (A) or posterior crossvein (PCV) phenotypes (B,C). D-H) 24hAPF control pupal wing (D) or wings expressing wild type *sog* (E) or *sogN* mutants (F-H), driven by shv-Gal4 and stained for pMad.

*sog* overexpression also results in PCV loss. We find that ubiquitous overexpression of *sogN* mutants in the wing (with MS1096-Gal4 driver) increases PCV loss when compared to *sogWT* overexpression (Fig. 4B). This difference is exacerbated when the overexpression is restricted to the longitudinal proveins by using the *shv*-Gal4 driver, with *sogN1* and *sogN2* again showing the strongest effects (Fig. 4C). These different effects on the PCV produced by local (ubiquitous, MS1096-Gal4) versus at-a-distance (longitudinal proveins, *shv*-Gal4) overexpression of *sog* mutants indicate that the transport of BMP signaling complexes in the extracellular space is affected in the *sogN1* and *sogN2* mutants.

Accordingly, overexpression of *sogWT* and *sogN* mutants in longitudinal proveins affects the pMad pattern in the pupal wing. In the longitudinal proveins, pMad staining is decreased and uneven in *sogWT* overexpression wings when compared to control wings (Fig. 4D,E) consistent with inhibition of BMP signaling by Sog in these domains. In contrast, the PCV appears enlarged and with increased pMad levels (Fig. 4E), indicating increased BMP transport from the longitudinal proveins to the PCV domain. Overexpression of *sogN* mutants seems to exacerbate these defects, with the PCV often appearing detached from the longitudinal proveins (Fig. 4F-H), consistent with the phenotypes observed in adult wings (Fig. 4A,C) and with increased BMP binding (Fig. 3D). Interestingly, the pMad pattern resulting from *sog* overexpression (decreased and non-uniform levels in longitudinal proveins and enlarged PCV) is reminiscent of the pMad pattern observed in *frc-* wings (compare Fig. 4F-H to Fig. 1D-E).

During embryogenesis, *sog* is expressed in the lateral neuroectoderm where it inhibits *dpp* from auto-activating (Biehs et al., 1996). Sog also diffuses dorsally to concentrate BMP activity to the dorsal-most region of the embryo (Mizutani et al., 2005, Shimmi et al., 2005, Ashe and Levine, 1999, Decotto and Ferguson, 2001). Peak BMP activity leads to dorsal expression of *zen*, *RACE* and *rho*, as well as the formation of the amnioserosa, which is absent in loss-of-function *sog* mutants (Ray et al., 1991, Yu et al., 2000). After gastrulation, amnioserosal cells can be detected with antibodies against the Kruppel (Kr) transcription factor. To assay the effects of *sogN* mutants in the embryo we drove early ubiquitous expression of *sogWT and sogN* mutants with a maternal Gal4 driver (matα-Gal4). *sogWT* overexpression does not significantly alter the domain of *zen* expression or the number of Kr+ cells, as previously shown (Yu et al., 2000) (Fig. 5A,B,F,G). However, overexpression of *sogN1* and *sogN3* reduces the domain of *zen* expression as well as the number of Kr+ cells (Fig. 5C,E,H,J,K,L), indicating that these mutants exert an inhibitory effect on BMP signaling.

**Figure 5.**
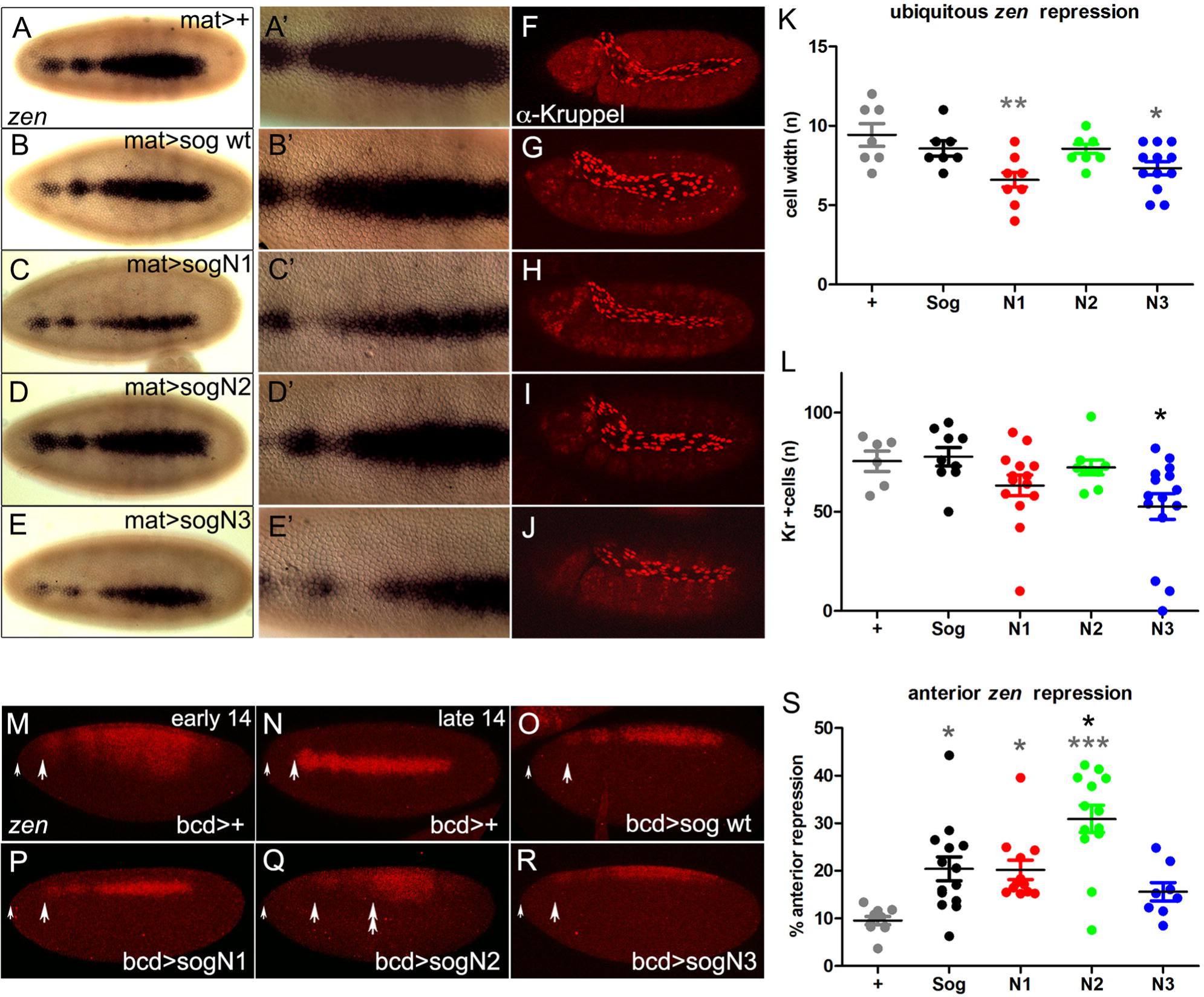
Loss of glycosylation enhances Sog function in the embryo. A-L) Embryonic pattern produced by expression of wild type *sog* or *sog* glycosylation mutants *sogN1, sogN2* or *sogN3*, with the ubiquitous matαGAL4 driver. Effects were analyzed by A-E) dorsal *zen* expression, quantified in K), or F-J) Kruppel protein in amnioserosal cells, quantified in L). M-S) Embryonic pattern produced by localized expression of wild type *sog* or *sog* glycosylation mutants *sogN1, sogN2* or *sogN3*, with the anteriorly restricted bcdGAL4/GCN driver. Effects were analyzed by dorsal *zen* expression. The domain of *zen* repression from the anterior tip of the embryo was measured and quantified in S). A small arrow points to the anterior tip of each embryo. Arrows in M-R point to the most anterior region of *zen* expression detected. Double arrow shows the region where *zen* is expanded relative to wild type *sog* by anterior *sogN2* expression (Q). Statistically significant differences based on Student’s t-test (***P≤0.001, **P≤0.01, *P≤0.05).

Taking into account that, in the wing, *sogN* mutants displayed different effects locally versus at-a-distance, we performed a similar analysis in the embryo. We induced localized expression of *sogN* mutants in the anterior tip of the embryo by using the *bcd-*Gal4 driver (*bcdGCN*-Gal4, (Yu et al., 2000), Fig. 5M-R). Anterior overexpression of *sogN1* and *sogN3* decreases the domain of *zen* expression to the same extent as *sogWT* (Fig. 5O,P,R). However, *sogN2* inhibits *zen* expression further away from the anterior tip (Fig. 5O,Q,S). In addition, in *bcd*>*sogN2* the *zen* domain is broad in the middle of the embryo, suggesting that SogN2 shuttles the active BMP complex towards the posterior end to a greater extent than wild type Sog (Fig. 5Q).

Altogether, the analysis of *sog* overexpression in the wing and embryo shows that the effects of individual mutations in putative Sog glycosylation sites vary depending on context, with SogN1 and SogN2 showing the greatest effects in the wing and SogN2 in the embryo. Nevertheless, it is important to point out that in all contexts above the loss of putative glycosylation sites increases the well-established effects of Sog to inhibit and/or shuttle BMPs.

### Glycan removal inhibits Sog endocytosis

Precise morphogen activity requires controlling the levels and spatial distribution of extracellular morphogens and their antagonists. In Drosophila, endocytosis regulates the amount of Sog protein in the extracellular space and thus BMP activity (Srinivasan et al., 2002, Negreiros et al., 2010). Sog suppresses the action of excess Gbb in the wing and Scw in the embryo, but not Dpp (Neul and Ferguson, 1998, Nguyen et al., 1998, Yu et al., 2000). Accordingly, extracellular Sog is able to bind the BMPs Scw and Gbb, and with lower affinity to Dpp. However, in the presence of Tsg, Sog binds to and antagonizes both Dpp heterodimers and homodimers (Ross et al., 2001, Shimmi et al., 2005). Importantly, Sog retrieval from the extracellular space also increases in the presence of Tsg (Fig. 6A), regardless of the BMP involved. Sog retrieval from the extracellular space is dependent on BMP and Integrin receptors (BMP receptor type I Tkv, βPS integrin receptor myospheroid, αPS1 integrin receptor multiple edematous wing and αPS2 receptor inflated), since *tkv*, *mys, mew* and *if* knockdowns decrease intracellular Sog amount in S2 cells (Fig. 6B). This indicates that Sog retrieval in the presence of Tsg requires an interaction with a Dpp homo or heterodimers and with Integrin receptors, following endocytosis of Sog and possibly BMPs that remain bound to the complex.

**Figure 6.**
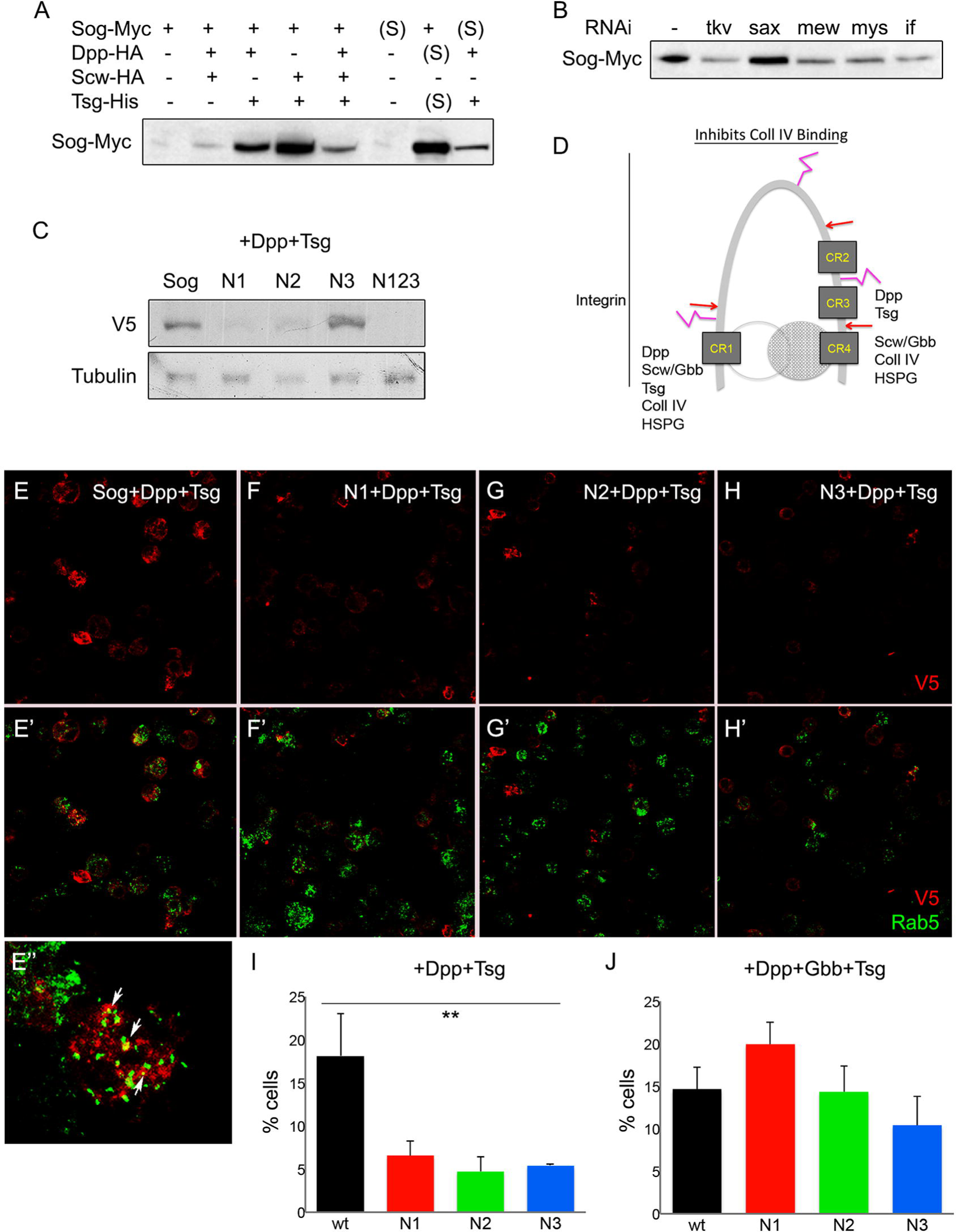
Sog levels are controlled by glycan and receptor-based retrieval from the extracellular space. A) S2 cells transfected with wild type *sog-myc* construct and different combinations of constructs producing Tsg and BMP ligands. Cellular amounts of Sog are detected with anti-myc antiserum. Sog amounts increase in the presence of Dpp plus Tsg. These cellular levels correspond to Sog retrieved from the medium, since equivalent intracellular amounts are observed when medium (S) from construct-expressing cells is added to cells that do not express *sog-myc.* B) Intracellular Sog levels retrieved from Sog-myc+Dpp-HA+Tsg-His medium decrease by knocking down the Dpp receptor *tkv* or αPS1 (*mew*), αPS2 (*if*) or βPS (*mys*) integrin expression. C) S2 cells transfected with wild type *sog-V5* or *sogN1-V5*, *sogN2-V5*, *sogN3-V5* or tripple *sogN123-V5* mutant constructs in the presence of constructs producing Tsg and Dpp. Sog retrieval decreases by mutating the N1 and N2 sites. D) Predicted conformation of Sog protein with regions depicted for binding to established extracellular partners. E-J) Medium from S2 cells transfected with *dpp-HA* and *tsg-His* plus wild type *sog-V5* or *sogN1-V5*, *sogN2-V5*, or *sogN3-V5*, was added to naive cells and imunolabelled with anti-V5 (red) and anti-Rab5 (green, E’-H). E’’) High magnification for wild type Sog-V5 and Rab5 shows partial co-localization, indicative of Sog endocytosis. These cells were quantified in (I), showing that loss of N1, N2 and N3 glycosylation sites decreases the number of cells with V5+Rab5+ endocytic punctae. J) In cells expressing *dpp-HA, gbb-His* and *tsg-His* plus wild type *sog-V5* or *sogN1-V5*, *sogN2-V5*, or *sogN3-V5,* the number of cells with V5+Rab5+ punctae is equivalent. Statistically significant differences based on Student’s t-test, **P≤0.01.

Loss of putative glycosylation sites decreases Sog retrieval in S2 cells (Fig. 6C). In the presence of Dpp and Tsg, SogWT-V5 is retrieved from the medium by cells that do not express V5-tagged Sog. In the same condition, intracellular levels of SogN1, SogN2 and the triple mutant SogN123 decrease compared to SogWT (Fig. 6C). This indicates that glycosylated residues at the Sog N-terminus and stem are required for an interaction with membrane receptors that retrieve Sog from the extracellular space. Furthermore, Sog retrieval from the extracellular space results in internal vesicle-bound Sog. Co-localization analyses shows that a significant amount of intracellular Sog positive punctae correspond to Rab5 positive endocytic vesicles (Fig. 6E). Notably, the percentage of cells displaying Sog associated to endocytic vesicles decreases by mutating any of the three putative glycosylation sites N1, N2 or N3 (Fig. 6F-H and 6I). Therefore, a likely interpretation of these results is that loss of Sog glycosylation leads to a decrease in receptor-based Sog endocytosis, thus increasing the amount of extracellular Sog. Interestingly, retrieval of wild type and mutant Sog is equivalent in the presence of Tsg+Dpp+Gbb (Fig. 6J). Thus, differential distribution of BMPs in the tissue relative to glycosylated Sog and Tsg could provide additional levels of complexity in the regulation of BMP activity by Sog endocytosis.

## Discussion

### N-linked glycosylation modulates Sog function

Here we have undertaken a biochemical and functional analysis of glycosylation in the BMP antagonist Sog. Our results confirm that Sog is glycosylated and show that three sites target for N-glycosylation are evolutionarily conserved. While appropriate glycosylation may be important for protein folding in general (Caramelo and Parodi, 2015, Jayaprakash and Surolia, 2017) and thus may impact the total amount of active glycoprotein produced, our results indicate that Sog glycosylation controls a higher level of Sog function. Our *in vivo* functional analysis shows that loss of conserved glycosylation sites enhances extracellular Sog activity in all the contexts tested. Specifically, mutating conserved glycosylation sites increases BMP antagonism in the wing (N1 and N2) and embryo (N1 and N3), consistent with an increase in Dpp binding in S2 cells (N1). Furthermore, loss of Sog glycosylation sites also increases the range of Sog effects in the pupal wing (N1 and N2) and embryo (N2), and decreases the retrieval of Sog protein from the extracellular space in S2 cells (N1 and N2, to a smaller extent N3). Therefore, our results consistently point to a role for the first glycosylation site (N1) in dampening BMP antagonism and for the first and second glycosylation sites (N1 and N2) in restricting BMP shuttling by Sog. Notably, loss of each individual Sog glycosylation site results in slightly variable effects depending on context, which may reflect spatial and temporal differences in interacting partners.

The presence of glycans on extracellular proteins frequently alters the degree and quality of interactions with the extracellular matrix and with other secreted proteins (Tauscher et al., 2016, Dennis, 2017, Jayaprakash and Surolia, 2017). By analyzing the position of the putative Sog glycosylation sites one might gain insight to explain how glycan addition may regulate Sog binding to interacting partners. It has been suggested that Chd and Sog assume a horseshoe-like conformation that enables cooperative BMP binding (Troilo et al., 2015, Larraín et al., 2000, Shimmi et al., 2005). In this arrangement, specific N- and C-terminal regions interact with BMPs (CR1, CR3 and CR4) (Sawala et al., 2012, Troilo et al., 2015), Tsg proteins (CR1 and surrounding) (Yu et al., 2004), heparan sulfate proteoglycans (HSPGs, CR1 and CR4) (Jasuja et al., 2004), integrin receptors (N-terminal) (Araujo et al., 2003, Larraín et al., 2000) and collagen (CR1 and CR4) (Sawala et al., 2012) (Fig. 6D). Taking into account that most glycans are present at the external face of Drosophila glycoproteins, it is unlikely that Sog glycosylation directly alters BMP dimer binding at internal sites (Jayaprakash and Surolia, 2017). Alternatively, glycosylation could modify Sog conformation, indirectly changing the interaction with BMPs. Another possibility is that Sog glycosylation provides the structural basis for interaction of the Sog/BMP complex with the extracellular matrix and transmembrane receptors. These interactions could secondarily affect BMP binding and metalloprotease cleavage dynamics as well as movement of the BMP complex in the tissue. Structural analysis of the Sog protein should allow testing this prediction in the future.

The most consistent results gathered in this manuscript concern the N1 and N2 sites. The N-terminal, Cystein Repeat 1 (CR1) containing Sog domain, harbors a Tld/Tlr site proximal to N1 (Marqués et al., 1997). Metalloprotease cleavage at this site generates an N-terminal fragment termed Supersog, which is a stronger BMP antagonist than full-length Sog (Yu et al., 2000), and is able to bind Dpp homodimers, unlike full-length Sog that only binds BMP heterodimers (Yu et al., 2000, Shimmi et al., 2005) (Fig. 3). Therefore, one mechanism by which SogN1 could increase BMP antagonism would be to favor the generation of Supersog fragments. However, our results contradict this interpretation, since loss of this first glycosylation site does not favor N-terminal Sog cleavage or modify the Sog cleavage pattern. Alternatively, loss of N1 glycosylation may modify interactions that take place near this site. For instance, by favoring BMP binding, either by increasing binding strength or decreasing the requirement for accessory proteins such as Tsg, ultimately resulting in less BMP available for receptor activation. This interpretation is consistent with the ability of SogN1 to bind Dpp in the absence of Tsg, unlike wild type Sog that requires either a BMP heterodimer for binding or the presence of Tsg to bind Dpp homodimers. Based on this observation, it is tempting to suggest that differential glycosylation at the N1 site could be used as a mechanism to alter the Tsg requirement for BMP binding, complex formation and shuttling. Also noteworthy is the observation that vertebrate Chd binds BMP4 in the absence of Tsg (Oelgeschläger et al., 2000). It will be interesting to investigate in the future whether the N1 site is equally glycosylated in different developmental contexts of the fly and in vertebrate Chordin homologs.

### Stem (N2) Sog glycosylation controls the BMP activity range

An essential and conserved feature of Sog and its vertebrate Chd homologs is the ability to form highly dynamic complexes with BMP, Tsg and Tld proteins. During embryonic dorsal-ventral patterning Sog and Chd form multiprotein complexes that shuttle BMPs, concentrating BMP activity away from the site of Sog or Chd synthesis in a source-and-sink model of morphogen gradient formation (Bier and De Robertis, 2015). This model relies on the ability of Sog and Chd to diffuse in the extracellular space (Srinivasan et al., 2002, Plouhinec et al., 2013) and to interact with BMP, Tsg and Tld proteins (Mizutani et al., 2005, Umulis et al., 2006, Plouhinec et al., 2013, Reversade and De Robertis, 2005, Ambrosio et al., 2008, Inomata et al., 2008). In the pupal wing, formation of similar Sog/BMP complexes and shuttling BMPs away from the Sog source is also important for PCV formation (Serpe et al, 2008).

By altering the N2 Sog glycosylation site we observe BMP inhibition in the embryo along a greater extension relative to wild type Sog. A slight increase in *zen* expression away from the source of anterior *sogN2* expression is also observed, indicating that BMPs are shuttled further way from the SogN2 source in the process (Figure 5Q). This effect is strongest in the wing, where *sogN2* expression in the provein territory leads to inhibition of PCV formation close to the source (generating detached PCV and pMAD staining) and increased pMAD staining away from the source (Fig. 4G). These results can be interpreted as indication that Sog glycosylation at the N2 site restricts Sog diffusion and consequently how far from the source it is able to shuttle BMPs. Remarkably, the N2 site is the best conserved among Chordins (Fig. S2B), indicating that the range of BMP activity might be regulated by Chd stem glycosylation in vertebrates as well.

Interestingly, in *frc*-wings Sog distribution is abnormal, with high amounts entering the provein domain, away from the intervein Sog-producing source. An uneven pattern of pMAD inside the provein territory is also observed in *frc-*pupal wings, akin to the effect of overexpressing *sogN* inside the provein. Therefore, despite the generalized decrease in glycosylation expected in *frc-*wings, if the altered pMAD pattern in *frc-*results in great part from unglycosylated Sog entering the provein domain, then an important function of Sog glycosylation would be to control the even spread of BMPs along the center of the provein domain, leading to the formation straight, continuous veins (Yu et al., 1996).

Other developmental processes that require the precise control of BMP spread have been reported to depend on glycan residues. In the female germline stem cell (GSC) niche, Dpp diffusion is restricted to one cell diameter in order to maintain the undifferentiated or GSC renewal state (Harris and Ashe, 2011). The collagen IV Viking and the proteoglycan Dally bind to extracellular Dpp near anterior GSCs and prevent it from spreading further to the posterior, allowing the differentiation of cystoblasts (Guo and Wang, 2009, Wang et al., 2008, Hayashi et al., 2009). Similarly, during embryonic dorsal closure, *mummy* is required to define a highly restricted BMP activity field (Humphreys et al., 2013). The *mummy* gene codes for a key enzyme that generates UDP-N-acetylglucosamine for the synthesis of O- and N-linked glycans, as well as glycolipids and GPI anchors (Schimmelpfeng et al., 2006, Tonning et al., 2006). Therefore, glycosylated Sog, Viking, Dally and Mummy may be classified as a novel type of BMP regulators that can transform BMP from a long-range to a short-range signal.

### Glycans modulate BMP activity range by controlling receptor-based Sog endocytosis

The resident time of a secreted morphogen in the extracellular millieu has a direct impact on the range it is able to exert its effects (Gonzalez-Gaitan and Jülicher, 2014). As a secreted morphogen, extracellular Sog levels are regulated by endocytosis. This has proven important for Sog spread during embryogenesis (Srinivasan et al., 2002) and wing development (Araujo et al., 2003, Negreiros et al., 2010). Importantly, Sog glycosylation may be required for endocytosis. Our results show that Sog retrieval from the extracellular space requires Dpp (Tkv) and αPSβPS Integrin (Mys, Mew and If) transmembrane receptors (Fig. 6B). Accordingly, intracellular Sog levels are low inside *mew-* and *mys-* clones, localized in the intervein domain of the dorsal wing epithelium. Therefore, αPS1βPS integrin receptors regulate intracellular Sog levels in the pupal wing (Negreiros et al., 2010). Our present results show that by mutating Sog glycosylation sites N1 and N2 the amount of Sog retrieved from the extracellular space reduces. These sites are located inside the region of the Sog molecule responsible for Integrin binding (Araujo et al., 2003). Altogether, these results suggest that glycan-based interactions with membrane receptors regulate how much Sog is internalized, thus controlling the amount of extracellular Sog capable of shuttling BMP complexes. Future biochemical analysis should allow evaluate whether glycans are required for a direct interaction between Sog and specific Integrin receptor subunits.

Interestingly, Integrin receptors bind Xenopus Chordin, controlling the amount of Chordin that is internalized (Larraín et al., 2000). Taking into account the structural similarities between Sog and Chd, and the presence of conserved glycosylation sites, the possibility that glycan-based Sog endocytosis is conserved should be investigated. On the other hand, in the Drosophila embryo, Integrins regulate Dpp signals downstream of the receptor (Sawala et al., 2015), unlike the interaction described with Sog in the wing. However, Sog endocytosis is important for dorsal diffusion of Sog-containing BMP complexes (Srinivasan et al., 2002), suggesting that receptor-mediated Sog endocytosis in the embryo does not involve Integrins.

### Addition of sugar side chains as a general mechanism to regulate BMP function

Several molecules that interact with BMP ligands are modified by glycans, and BMPs themselves are subject to glycan addition (Hang et al., 2014, Lowery et al., 2014, Tauscher et al., 2016). Among these molecules are proteoglycans (Guo and Wang, 2009, Häcker et al., 2005, Selleck, 2000), Tsg family proteins (Billington et al., 2011), as well as Sog/Chd (Marqués et al., 1997). Therefore, disorders of glycosylation or changes in glycan availability will have a compounded impact on BMP function that increases in complexity as increases the number of glycosylated partners in a given biological context. While this a complex matter that shall not be resolved without mathematical modeling, examining the effects of individual glycosylation events provides us with the essential information to start building such models. Moreover, to examine protein glycosylation in light of evolution hints to the importance of glycan site acquisition.

Here we have shown that Sog function is regulated at specific, evolutionarily conserved glycosylation sites. We observed that mutating these sites increases BMP antagonism and/or BMP complex diffusion. The slight variability of effects observed in the different contexts is in consonance with the fact that the population of sugars attached to each glycosylated asparagine in a mature glycoprotein depends on the cell type, physiological and metabolic status of the cell in which the glycoprotein is expressed (Dennis et al., 2009; Dennis JW 2017). Therefore, altering the Sog glycosylation status during development could add great complexity to regulation of Sog glycoprotein function and consequently BMP activity.

## Materials and methods

Lines used in this study were: loss-of-function *frc*[00073], crossed into a TM6-Tb balancer to identify homozygous pupae, *dpp*[shv], and the MS1096- and matα Gal4 drivers, obtained from the Bloomington Indiana Stock Center. The shv-GAL4 driver was a gift of José DeCelis. All drivers have been described previously (Yu et al., 1996, Sotillos and De Celis, 2005).

### Constructs for S2 cell expression

A *Drosophila melanogster sog* cDNA cloned into pBluescript (Yu et al., 1996) was mutated at codons 179 (N1), 520 (N2), or 821 (N3), replacing Asn for Gln (CAA) (Retrogen, USA). Production of double (N23) mutated construct was performed by pBS-SogN1 and pBS-SogN2 digestion with SmaI/Xba and the triple (N123) mutated construct was performed by pBS-SogN1 and pBS-SogN23 digestion with SmaI/NotI. Wild type and mutant *sog* sequences were amplified and inserted into KpnI/NotI sites in pMT-V5/HisC (Invitrogen) to produce C-terminal tagged inducible constructs.

### Transgenic constructs

Constructs for Gal4/UAS mediated expression in flies were generated in pTiger, a derivative of pUASp (Ferguson et al., 2012). For the pTiger-SogWT construct, pBS-SogWT and pTiger were digested with KpnI and XbaI and ligated using T4 DNA ligase (NEB). For the pTiger-SogN1, N2 and N3 constructs, SogN1, N2 and N3 sequences were amplified by PCR from pBS-SogN1, N2 and N3, digested with KpnI and XbaI and introduced into KpnI/XbaI digested pTiger using T4 DNA ligase (NEB). pTiger-Sog constructs were sent to Bestgene for injection and integrated into the *attP*VK00027-docking site using the ΦC31 system (Markstein et al., 2008). At least two lines for each construct were generated and presented consistent effects.

All constructs were sequenced before use in cell transfection or transformation. Further details and primers used for production of S2 cell and transgenic constructs are available upon request.

### S2 cell culture and transfection

S2 cells were cultured in Schneider’s cell medium (GIBCO, Carlsbad, CA) containing 10% heat inactivated fetal bovine serum. Cells were transfected with CellFectin (Invitrogen) according to manufacturer’s instructions. For pMT-sog-V5/His constructs protein expression was induced by treatment of 0.7 mM CuSO_4_ 24 h after transfection. Sog protein was harvested from medium 48h after CuSO_4_ treatment. Vectors for sog-myc and tsg-His expression are described in (Yu et al., 2000). tld-HA, dpp-HA (Marqués et al., 1997), gbb-FLAG and Mad-FLAG (Ross et al., 2001) were a generous gift from M. O’Connor. Transfection product proteins were produced in the absence of glycosylation by incubating with tunicamycin (10ug/ml). RNA interference for *mys*, *mew*, *if*, *sax* and *tkv* was performed as in Kang and Bier 2010, with double stranded RNA synthesized from primers listed in Supplementary Material.

### Elisa

96-well plates (Nunc) were sensitized with 100 μl/well (4μg/ml) of anti-His, the capture antibodie, overnight at 4°C. Wells were washed 3 times with PBS pH 7.2; Tween-20 0.025% (PBST) and blocked with PBS supplemented with 10% FBS (PBS/10% SFB) in a volume of 200 μl/well. Plates were allowed to stand for 2 h at 37°C and washed 3 times with PBST. Subsequently, 50 μl of supernatants from S2 cell cultures were added to the wells. Plates were covered and incubated overnight at 4°C. After washing as plates 4 times with PBST, 100μl (4 μg/ml) of the detection antibodies (anti-V5, Invitrogen) was added to the wells. Plates were incubated for 1h at room temperature and washed 6 times with PBST. Subsequently, 100μl of streptoavidin-alkaline phosphatase (1μg/ml) diluted in PBS supplemented with 10% FBS were added to each well. Plates were incubated for 3h at room temperature. Subsequently, as plates were washed 8 times with PBST and so on in each well, 100μl of the 1.0mg/mL solution substrate bis-azine ethyl benzothiazole sulfonic acid (Sigma) diluted in 20 mM Tris, 100 mM MgCl_2_. Readings were performed on the Beckman Coulter AD340 reader with 405nm filter.

### Immunoblotting and immunoprecipitation

Medium or cell lysates from transfected S2 cells collected 72h after transfection. Cells were harvested and lysed with 200 Ll ice cold lysis buffer (50 mM Tris-HCl pH 7.5, 150 mM NaCl, 0.1% NP-40, protease inhibitor cocktail - Complete, Behringer Manheim) and centrifuged for 20 min at 4 °C to remove cell debris. After preclearing of total cell lysates with protein-G Sepharose beads for 20 min, the lysates were incubated with 30 μl of antibody bound agarose beads (crosslinked with monoclonal anti-V5 antibody, Invitrogen) at 4 °C for 3 h. Beads were washed 3 times with lysis buffer, then boiled in SDS-PAGE sample buffer at 95 °C for 5 min, and electrophoresed on 10% Bis-Tris gel and detected by immunoblotting onto PVDF membranes using anti-V5 (1:2000).

### Immunofluorescence and in situ hybridization in pupal wings and embryos

24-48h pupal wings were fixed for 10 min in 4% PFA in PBS, pH 7.0 and permeabilized in PBST buffer (PBS + 0.1% Tween20), blocked with 5% normal goat serum, incubated with primary antibody overnight at 4°C (rabbit anti-phospho-Smad1/5, Cell Signalling 41D10, 1:500; mouse anti-βIntegrin 1:500 DSHB; anti-Sog 8A or 8B antisera 1:500) and subsequently incubated with Alexa fluor secondary antibodies. Embryos were fixed and devitelinized in methanol using standard proceedures, after dechorionation in bleach. Embryos were blocked with 5% normal goat serum in PBST for 1 h and incubated with the primary anti-Kr antibodies (1: 1000, kind donation of Francisco Lopes) overnight at 4 °C. Secondary antibodies used were Alexa Fluor 488 goat anti-rabbit and Alexa Fluor 568 goat anti-mouse (1:500; ThermoFisher Scientific, USA), incubated for 1 h at room temperature. Hoechst 33342 was used at 1 □g/ml for nuclear stains. The fluorescence was detected by confocal microscopy using a Leica SP5-AOBS microscope (Leica Microsystems, Wetzlar, Germany). In situ hybridization was performed as previously described (Araujo and Bier, 2000).

### Protein sequence analysis

Short gastrulation homolog protein sequences were downloaded from Flybase or NCBI for the 12 *Drosophila* sequenced genomes, using the 2015 release. Sequence accession numbers are listed in Supplemental Material. Protein sequences were aligned using ClustalW in the MacVector (MacVector, Cary, NC, USA) program. Glycosylation sites were predicted in each protein using NetNGlyc 1.0.

### Rab5/Sog co-localization analysis

Sog endocytosis was analyzed by adding medium from control, wild-type Sog or N mutant Sog expressing cells, grown in the presence of Dpp-HA and Tsg-His, to non-transfected cells. Rab5 and Sog (wild type or mutant) were detected in the receiving cells with rabbit anti-Rab5 (1:500, Abcam, UK) and mouse anti-V5 (1:500) antisera and subsequently incubated with Alexa fluor secondary antibodies. Co-localization was analyzed with Image J software in single confocal optical sections. The number of cells with at least 2 Rab5+V5+ punctae was quantified for each condition.

## Acknowledgements

We are indebted to Mihaela Serpe and Michael O’Connor for kindly providing BMP pathway constructs and flies. We thank Attilio Pane and members of the Araujo and Todeschini labs for suggestions and critical reading of the manuscript. Monoclonal antibodies against Integrin subunits were obtained from the Developmental Studies Hybridoma Bank, created by the NICHD of the NIH and maintained at The University of Iowa, Department of Biology, Iowa City, IA 52242. Stocks obtained from the Bloomington Drosophila Stock Center (NIH P40OD018537) were used in this study.

## Author contributions

EN, SH, KY, AC, KC and HA performed experiments. All authors analyzed the data. AT, EB and HA developed the approach. SH, KC, EB, AT and HA prepared or edited the manuscript.

